# The C99 domain of the amyloid precursor protein is a disordered membrane phase-preferring protein

**DOI:** 10.1101/2020.11.25.397893

**Authors:** Ricardo Capone, Ajit Tiwari, Arina Hadziselimovic, Yelena Peskova, James M. Hutchison, Charles R. Sanders, Anne K. Kenworthy

## Abstract

Processing of the amyloid precursor protein (APP) via the amyloidogenic pathway is associated with the etiology of Alzheimer’s disease. The cleavage of APP by β-secretase to generate the transmembrane 99-residue C-terminal fragment (C99) and subsequent processing of C99 by γ-secretase to yield amyloid-β (Aβ) peptides are essential steps in this pathway. Biochemical evidence suggests amyloidogenic processing of C99 occurs in cholesterol- and sphingolipid-enriched liquid ordered phase membrane raft domains. However, direct evidence that C99 preferentially associates with rafts has remained elusive. Here, we test this idea by quantifying the affinity of C99-GFP for raft domains in cell-derived giant plasma membrane vesicles. We find that C99 is essentially excluded from ordered domains in HeLa cells, SH-SY5Y cells and neurons, instead exhibiting a strong (roughly 90%) affinity for disordered domains. The strong association of C99 with disordered domains occurs independently of its cholesterol binding activity, homodimerization, or the familial Alzheimer disease Arctic mutation. Finally, we confirm previous studies suggesting that C99 is processed in the plasma membrane by α-secretase, in addition to the well-known γ-secretase. These findings suggest that C99 itself lacks an intrinsic affinity for raft domains, implying either that amyloidogenic processing of the protein occurs in disordered regions of the membrane, that processing involves a marginal sub-population of C99 found in rafts, or that as-yet-unidentified protein-protein interactions involving C99 in living cells drive it into rafts to promote its cleavage therein.

## Introduction

Alzheimer’s disease is a progressive neurodegenerative disorder causing dementia that afflicts around 40−50 million persons worldwide (1). The amyloid precursor protein (APP), one of the key transmembrane proteins associated with Alzheimer’s disease, undergoes a complex and competing series of proteolytic processing events (2–5). APP is predominantly processed via the non-amyloidogenic pathway, in which APP is first cleaved by α-secretase in the middle of the amyloid forming sequence to release an 83 residue C-terminal fragment (C83), followed by cleavage of C83 in its transmembrane domain by γ -secretase. The amyloidogenic pathway involves sequential cleavage of APP first by β-secretase, also known as β-site amyloid precursor protein cleaving enzyme 1 (BACE1). BACE1 cleaves APP to generate a 99 amino acid long transmembrane C-terminal fragment described as APP-CTFβ or C99. C99 is then processively cleaved by γ-secretase (6,7), ultimately generating 40-mer Aβ40 and 42-mer Aβ42 peptides in addition to several less abundant species (8,9). The Aβ peptides aggregate and form oligomers that trigger a series of events resulting in synaptic dysfunction and, ultimately, neuron cell death causing progressive memory loss in Alzheimer’s patients. Aβ oligomers go on to aggregate and seed the amyloid plaque deposits that are detected in the brain of Alzheimer’s disease patients (10–12).

A variety of evidence suggests that amyloidogenic processing of APP occurs in lipid (membrane) raft domains (13–18). Current models suggest rafts are dynamic cholesterol and sphingolipid-enriched nanoscopic membrane domains that share features of ideal liquid ordered (Lo) membrane domains and function to compartmentalize cellular processes [for recent reviews, see (19–21)]. Early biochemical studies employing detergent-resistant membranes (DRMs) as a measure of raft association suggested that a fraction of APP and its C-terminal fragments, along with β-secretase and γsecretase enzymes, are localized within lipid raft fractions to varying degrees (22–35). Studies using cholesterol depletion to compromise the integrity of lipid rafts have shown that β-secretase mediated processing of APP is reduced in absence of cholesterol, leading to the suggestion that the amyloidogenic pathway is cholesterol dependent and thus raft dependent as well (13,36–38). The final proteolytic event of the amyloidogenic pathway, cleavage of C99 by γ-secretase to release Aβ, also has been proposed to be raft dependent (39,40). This model is based in part on observations that the C99 C-terminal fragment of APP is present in detergent insoluble or detergent resistant membranes in some cell types, which also led to the suggestion they may contain raft targeting traits (28,34). However, it has long been argued that DRMs cannot be directly equated with lipid rafts (19,41,42). Cholesterol depletion is also an inconclusive test for whether a particular process is raft associated, as the methods use do not necessarily remove only cholesterol from raft domains, and are known to affect a variety of cellular pathways and properties of cell membranes (43,44). Studying rafts in living cells also remains an ongoing challenge (19–21). Thus, complementary approaches are needed to establish the localization of the components of the amyloidogenic pathway within distinct membrane domains and to illuminate possible roles for rafts in Alzheimer’s disease.

Biophysical studies from our group have shown cholesterol directly binds to C99 (18,45), raising the intriguing possibility that this could serve as a signal that enables C99 to associate with lipid rafts. To test this idea, in previous work we investigated the preferred localization of C99 in raft versus non-raft phases in synthetic giant unilamellar vesicles (GUVs) generated from a ternary lipid mixture that supports the formation of co-existing liquid-ordered (Lo) and liquid disordered (Ld) domains (46). The results of those studies showed that purified C99 is predominantly localized to non-raft Ld domains in this system. More recently, the phase preference of a γ-secretase substrate consisting of the C-terminal 76 residues of APP for Lo versus Ld domains was evaluated following its direct insertion into supported lipid bilayers using atomic force microscopy (47). Interestingly, the substrate was found to associate with both ordered and disordered domains, with a preference for ordered domains. However, it is unclear if these studies report on the phase preference of C99 itself since this construct is 23 residues shorter than C99. In addition, both of these studies were carried out in model membrane systems lacking the full complexity of biological membranes. Whether C99 has an intrinsic affinity for associating with ordered domains in cell membranes thus remains an open question.

To further investigate this issue, we here utilize giant plasma membrane vesicles (GPMVs) as a model. Derived from the plasma membrane of live cells, these μm-sized vesicles have served as a useful model system to investigate mechanisms that regulate membrane phase behavior and control the localization of lipids and membrane-associated proteins with raft versus non-raft domains (48–54). Similar to biomimetic model membranes such as GUVs, GPMVs can undergo fluid-fluid phase separation into two coexisting domains: a raft-like ordered (Lo-like) phase and non-raft disordered (Ld-like) phase. These domains can sort exogenous as well as native protein and lipids and can be readily visualized and analyzed using conventional fluorescence microscopy techniques (48–51). Typically, such measurements are carried out at room temperature or lower, where the domains are microns in size. Importantly however, even at temperatures above the miscibility transition temperature where GPMVs appear by fluorescence microscopy to consist of a single membrane phase, the presence of nanoscopic domains can be detected (55,56). Furthermore, GPMVs maintain compositional complexity that is similar to native cellular membranes (48,51,52). GPMVs thus better reflect the complex environment of cell membranes and represent a more physiologically relevant model to study the localization or association of proteins with raft-like ordered domains versus disordered domains.

To establish the membrane domain phase preference of C99 in a complex cellular membrane environment, we studied GFP-tagged C99 in GPMVs derived from HeLa cells and the neuroblastoma cell line SH-SY5Y as well as in its differentiated neurons. We show that overexpressed C99-GFP has a strong preference for the disordered phase of GPMVs. Moreover, the high preference of C99 for the disordered phase is not only independent of cell type but also independent of C99 dimerization, does not require an intact cholesterol binding domain, and is unchanged in a disease-associated mutant form of C99 that contains the Artic E693G mutation. These findings suggest that C99 lacks an intrinsic affinity for raft domains and that localization of C99 to ordered domains may not be required for further processing along the amyloidogenic pathway.

## Results

### Treatment of cells with γ-secretase inhibitor enables visualization of C99-GFP in GPMV membranes

To determine whether C99 preferentially resides within raft-like ordered or non-raft disordered domains in GPMVs, we initially utilized HeLa cells, in which APP and C99 have commonly been studied (57,58). In addition, a significant fraction of HeLa cell-derived GPMVs are known to contain micron-scale coexisting ordered (raft-like) and disordered (non-raft) phases at room temperature (59–61), making them an excellent model to quantify the phase preference of specific proteins and lipids. The degree of raft association of a given protein within GPMVs is typically assessed by comparing its localization to that of ordered or disordered phase-specific marker dyes (48,49,52,62).

To visualize C99, we utilized a version of C99 tagged with EGFP on its C-terminus (C99-EGFP) that has been shown to be correctly recognized by γ-secretase as a substrate (57). When C99-EGFP was overexpressed in HeLa cells, the protein accumulated in the perinuclear region and was mainly excluded from the plasma membrane **(Supporting Figure S1A)**. In contrast, upon treatment of HeLa cells with the γ-secretase inhibitor DAPT for 24 h, some C99-EGFP accumulated on the plasma membrane **(Supporting Figure S1B)**, in agreement with previous studies (63,64).

Next, we generated GPMVs from HeLa cells transiently overexpressing C99-EGFP. The transfected cells were either treated with vehicle (DMSO) or with 20 μM DAPT for 24 h prior to the preparation of GPMVs, and GPMVs were generated in the continued presence of the inhibitors. The isolated GPMVs were subsequently labeled with DiD, a lipid-based dye and a known disordered phase marker (53,65) and then analyzed using confocal microscopy. Consistent with the absence of C99 in the plasma membrane in intact cells under control conditions, we were unable to detect C99 in the membrane of GPMVs prepared from HeLa cells in the presence of DMSO **(Figure 1A, E)**. Instead, the GFP signal was concentrated in the lumen of the GPMVs **(Figure 1A, E).** This suggests that C99-EGFP is proteolytically processed to release a soluble cytoplasmic fragment containing GFP prior to and/or during the preparation of the GPMVs. In contrast, in DAPT-treated GPMVs, we also observed membrane-associated fluorescence signal, indicating that at least a fraction of the C99-EGFP cleavage seen in non-treated cells is due to γ-secretase **(Figure 1B)**.

**Figure 1.**
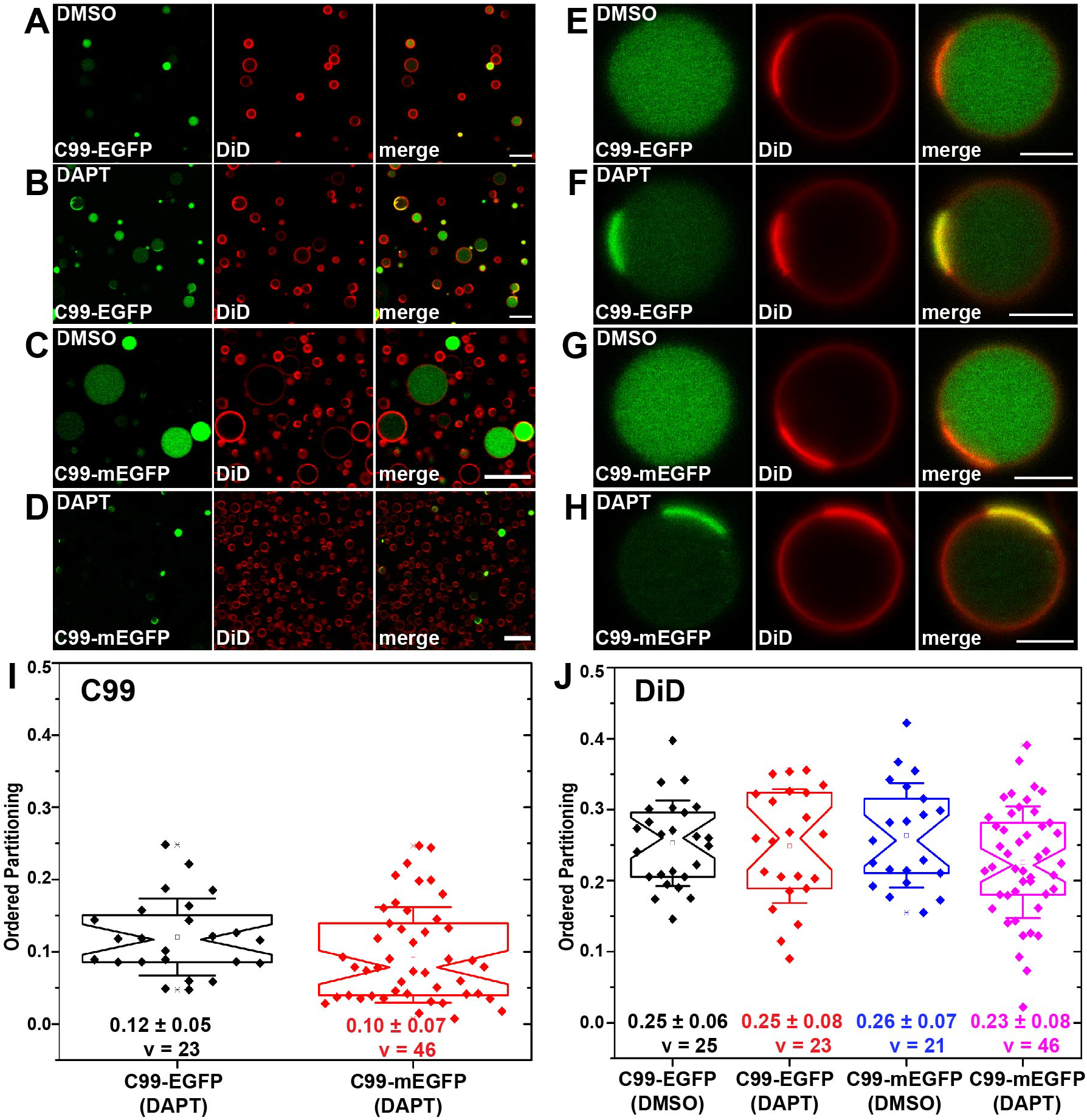
C99-EGFP and C99-mEGFP are excluded from the membrane of GPMVs under control conditions but accumulate in disordered membrane domains in response to γ-secretase inhibitor (DAPT) treatment. GPMVs were prepared from HeLa cells transfected with C99-EGFP or C99-mEGFP and treated with either DMSO or DAPT for 24 h prior to and during GPMV preparation. **(A-D)** Representative panorama images of GPMVs containing C99-EGFP (A, B) or C99-mEGFP (C, D) that were treated with either DMSO or DAPT as indicated on the panels. Scale bars, 20 μm. **(E-H)** Representative images of individual GPMVs treated with either DMSO or DAPT derived from cells expressing C99-EGFP (E, F) or C99-mEGFP (G, H). Scale bars, 5 μm. In A-H, the disordered phase marker (DiD) is shown in red, and GFP fluorescence is shown in green. **(I)** Quantification of ordered domain partitioning for C99 in DAPT-treated samples. **(J)** Quantification of ordered domain partitioning for DiD for samples treated with either DMSO or DAPT, as indicated. P_ordered_ values were calculated from 2-3 independent experiments with >10 GPMVs analyzed per replicate. Each data point represents a P_ordered_ measurement from a single GPMV. The mean ± SD is reported on the graph. v = number of vesicles analyzed.

Analysis of phase-separated GPMVs from DAPT-treated cells that were co-stained with the disordered phase marker DiD revealed that C99-EGFP colocalized with DiD, and hence preferentially resided in the disordered phase **(Figure 1F)**. Essentially no GFP signal could be detected in ordered domains, indicating the protein has a very strong preference for disordered domains **(Figure 1F)**. This was verified across several independent experiments in which we quantified the ordered phase preference of C99-EGFP in multiple GPMVs by calculating the ordered domain partitioning fraction *P*_*ordered*_ as described in the Materials and Methods (**Supporting Figure S2)**. In this analysis, a *P*_*ordered*_ between 0 and 0.5 means the protein prefers the non-raft phase, whereas *P*_*ordered*_ between 0.5 and 1.0 is indicative of a raft-preferring protein. *P*_*ordered*_ for C99-EGFP was strongly non-raft preferring, 0.12 ± 0.05 (Fig. 1I, v = 23 vesicles). In comparison, the non-raft domain marker DiD itself had a *P*_*ordered*_ of ~0.25 both in vehicle-treated GPMVs and in DAPT-treated GPMVs (**Figure 1J**).

We noticed that in GPMVs treated with DAPT some GFP fluorescence could still be observed in the lumen, despite using the inhibitor at concentrations above those previously reported to yield maximal protection of EGFP-C99 from cleavage by γ-secretase (57). The presence of luminal GFP in GPMVs even when the γ-secretase inhibitor DAPT was present was unexpected since γ-secretase inhibition should in principle prevent processing of C99 to liberate the soluble fragments. We therefore considered the possibility that this may be related to properties of the C99-EGFP construct itself. Inspection of the construct revealed a Kozak sequence upstream of the GFP moiety. To rule out the possibility that free GFP was being expressed without C99 we deleted the Kozak sequence and the first methionine of EGFP. We simultaneously mutated alanine at 206 to lysine (A206K) to generate mEGFP to prevent possible EGFP-induced dimerization (66). We refer to this optimized version as C99-mEGFP in subsequent experiments.

Like C99-EGFP, C99-mEGFP was localized primarily in the perinuclear region in HeLa cells under control conditions and became enriched in the plasma membrane in response to DAPT treatment (**Supporting Figure S1**). The fluorescence signal for C99-mEGFP was again found exclusively in the lumen of GPMVs derived from cells treated with DMSO (**Figure 1C, G**), and partially localized to the membrane for GMPVs from cells treated with DAPT where it was strongly excluded from ordered domains, similar to the behavior of C99-EGFP (**Figure 1D, H, I**). These findings further support the notion that C99 prefers to localize to disordered domains over ordered domains.

The finding that some luminal GFP signal was still present in GPMVs derived from DAPT-treated cells expressing C99-mEGFP suggests the protein undergoes additional processing by proteases other than γ-secretase. We next investigated potential sources of this additional proteolytic activity. Since both C99-EGFP and C99-mEGFP yielded essentially the same results, all further experiments were carried out with C99-mEGFP.

### C99-mEGFP is processed by α-secretase and caspases in addition to γ-secretase in HeLa cells

In addition to being a substrate for γ-secretase, C99 can also be proteolytically cleaved by several other proteases. Both full length APP and C99 are substrates for caspases, and caspase cleavage of their cytosolic domain releases a soluble C31 fragment (67,68) [reviewed in (69,70)]. Notably, C99-EGFP has also been reported to undergo caspase cleavage in HeLa cells (57). C99 is also a substrate for α-secretase, leading to its conversion to C83 under conditions where γ-secretase is inhibited (58,71). We hypothesized that one or both of these activities may be contributing to the processing of C99-mEGFP.

To establish which proteolytic activities are operative on C99-mEGFP, we performed a series of experiments in which C99-mEGFP transfected cells were treated for 24 h with DAPT in various combinations with the cell permeable pan-caspase inhibitor z-VAD and the ADAM10 specific inhibitor GI254023X. The cells were then harvested for western blotting and whole cell extracts were probed with antibodies using an N-terminal specific antibody for Aβ/C99, anti-GFP antibodies, and an antibody against the C-terminal portion of C99 (**Figure 2 and Supporting Figures S3 and S4**).

**Figure 2.**
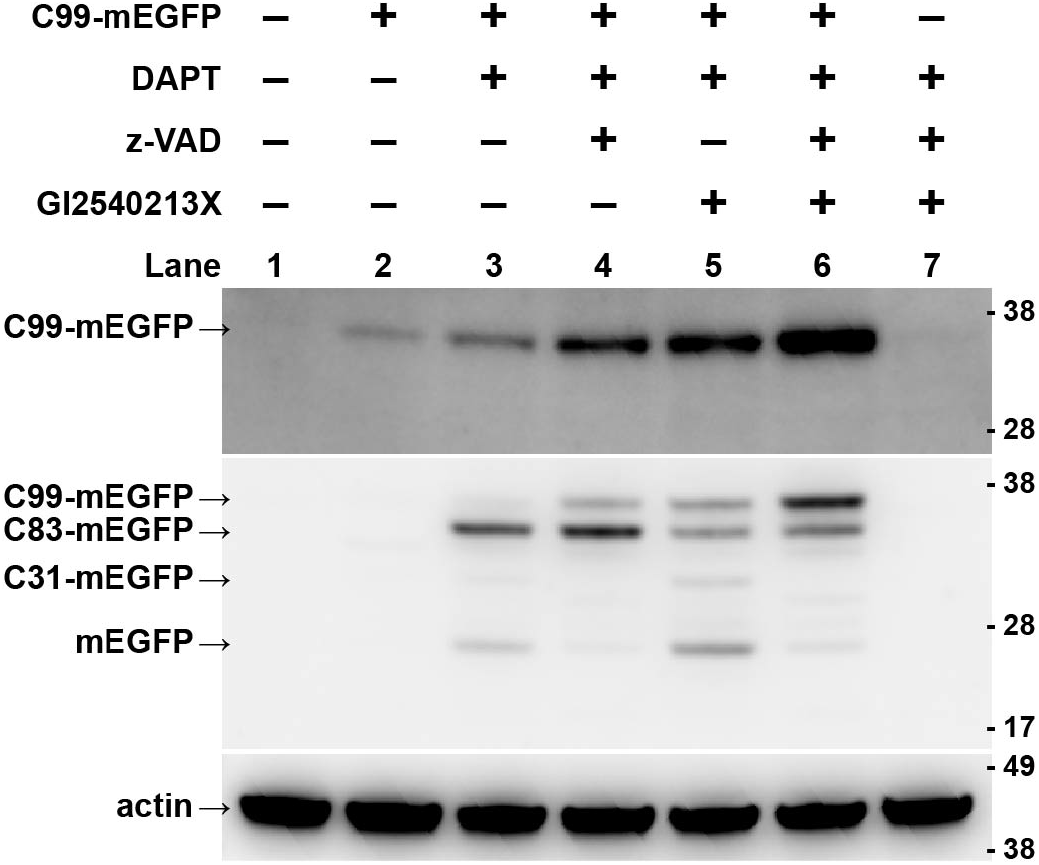
C99-mEGFP is cleaved by α-secretase and caspases in addition to γ-secretase. HeLa cells were either mock transfected or transfected with C99-mEGFP and treated with either DMSO or the indicated inhibitors for 24 h prior to processing for western blotting. Inhibitors were used at the following concentrations: DAPT (20μM), GI254023X (20μM), and z-VAD (200μM). The final concentration of DMSO in all samples was 0.4%. 10 μg of protein was loaded in each lane. Blots were probed with an N-terminus specific anti-C99 antibody (mAb 82E1; top panel), anti-GFP (mAb JL-8; middle panel), or anti-actin (mAb MCA-5J11; bottom panel) as a loading control. Supporting Figure S3 shows the same protein samples probed with anti APP-CT20 and a different anti-GFP antibody. Additionally, images of the complete membranes and total protein staining are available as Supporting Figure S4.

C99-mEGFP has a predicted molecular weight of ~38 kDa. Under most conditions tested, we detected a band at this molecular weight using an N-terminally specific anti-C99 antibody, suggesting that at least a fraction of C99-mEGFP remained unprocessed in cells. In contrast, when probed using an anti-GFP antibody, a variety of cleavage products could also be observed. In the absence of any inhibitors, only a very faint band of intact C99-mEGFP could be seen in cell extracts by western blot (**Figure 2**, lane 2), possibly corresponding to intracellular membrane-associated C99 that is excluded from the GPMV preparation. In cells treated with DAPT (lane 3) an increase in intensity of the band corresponding to intact C99-mEGFP could be seen when compared to lane 2 (**Figure 2**, lane 3). However, we also observed bands running at the expected positions of C83-mEGFP and mEGFP, indicating that additional proteolytic processing of the protein was also occurring.

In the presence of a combination of DAPT and the pan-caspase inhibitor z-VAD (lane 4), we observed two main bands, one corresponding to intact C99-mEGFP and a second, stronger band running at the position expected for C83-mEGFP (**Figure 2**, lane 4). When a combination of the γ-secretase inhibitor DAPT and the α-secretase inhibitor GI2540213X was used, a major band corresponding to full length C99-mEGFP, together with weaker bands running at the position of C83-mEGFP and C31-mEGFP were obtained (**Figure 2**, lane 5). Finally, in the presence of all three inhibitors (hereafter referred to as triple inhibitor cocktail), the intensity of the C99-mEGFP band increased compared to lane 2 (**Figure 2**, lane 6) and the intensity of the mEGFP band was diminished. Thus, addition of triple inhibitor cocktail strongly increased the amount of C99-mEGFP and decreased the levels of C31- and C83-mEGFP-like bands.

Taken together, these results show that in the presence of DAPT, overexpressed C99-mEGFP is mostly processed both by α-secretase (to produce the C83-mEGFP protein) and by caspases (**Figure 2**, lane 3). This implies that in GPMVs derived from DAPT-treated cells (**Figure 1**), the membrane-associated form of the protein present is predominantly the C83 form, and the luminal GFP signal likely consists of a mixture of C31-EGFP, AICD-EGFP, and fluorescent EGFP fragments. Because the highest level of intact C99-mEGFP was seen in cells treated with the triple inhibitor cocktail, this cocktail was used in subsequent studies to preserve C99-mEGFP in the membrane so that its phase preference could be more directly investigated.

### C99-mEGFP localizes in the disordered phase of HeLa GPMVs derived in the presence of γ-secretase, α-secretase, and caspase inhibitors

We carried out studies in which we jointly inhibited γ-secretase, α-secretase and caspases prior to and during GPMV preparation in order to preserve the C99 form of the protein at the plasma membrane. In HeLa cells incubated with the triple inhibitor cocktail, C99-mEGFP localized as expected to the plasma membrane (**Figure S1 E**). We observed that GPMVs could be successfully isolated in the presence of the triple inhibitor cocktail, and that a significant fraction were phase-separated at room temperature (**Figure 3A**). In the presence of the triple inhibitors, C99-mEGFP again strongly co-localized with the disordered phase marker DiD and was essentially excluded from ordered domains, with a *P*_*ordered*_ of 0.13 ± 0.09 (**Figure 3B, C**). This was similar to the value obtained in GPMVs derived from cells treated with DAPT alone (**Figure 1**). Thus, over 85% of both C83-mEGFP (**Figure 1**) and C99-mEGFP (**Figure 3)** localize into the disordered membrane environment.

**Figure 3.**
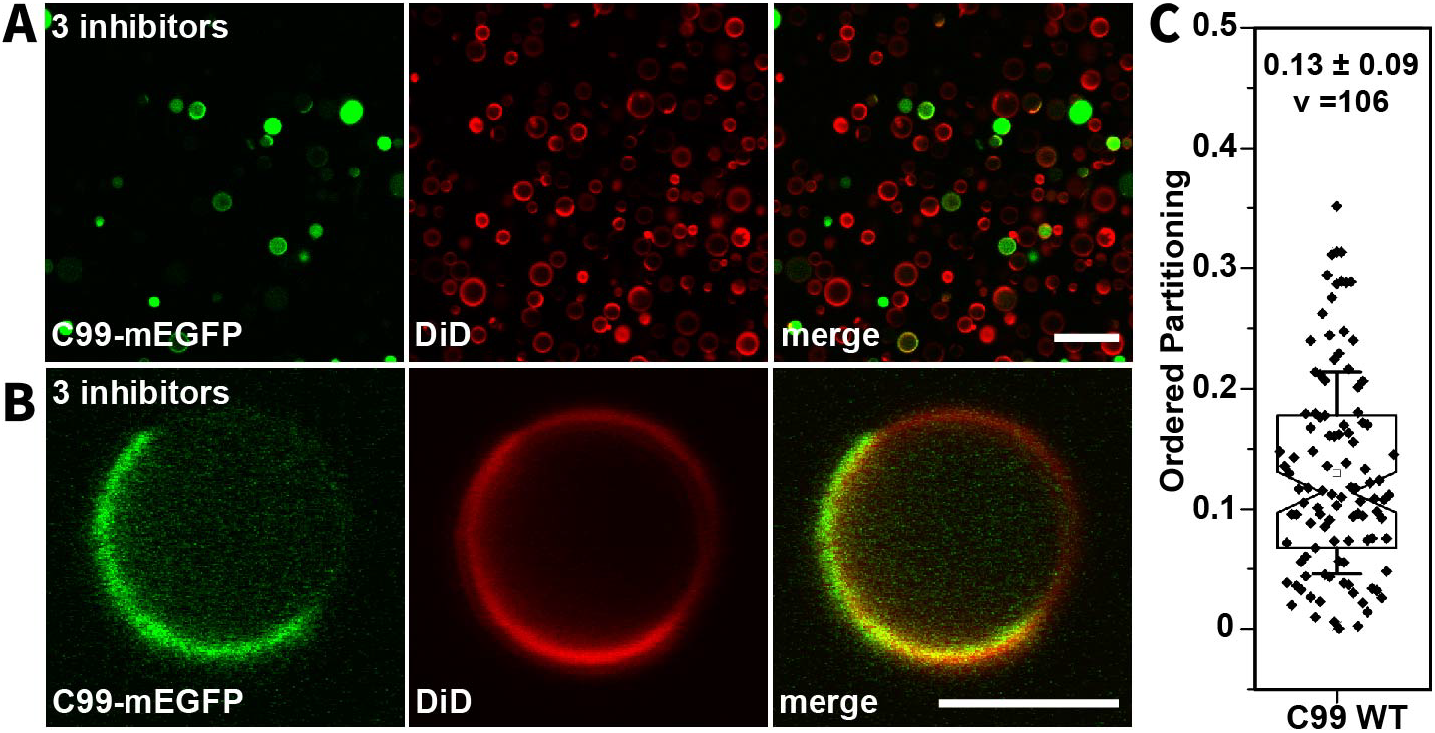
C99-mEGFP shows strong disordered-phase partitioning preference in GPMVs prepared in the presence of a mixture of γ-secretase, α-secretase, and caspase inhibitors. GPMVs were prepared from HeLa cells expressing C99-mEGFP subjected to treatment with a triple inhibitor cocktail as described in the Materials and Methods. **(A)** Representative image of a field containing multiple GPMVs. Note that some GPMVs exhibit GFP fluorescence primarily in the membrane while in others substantial fluorescence is observed in the GPMV lumen. Scale bar, 20 μm. **(B)** Representative image of an individual GPMV demonstrating colocalization of C99-mEGFP (green) with the DiD-positive disordered phase (red). Scale bar, 5 μm. **(C)** Quantification of ordered domain partitioning for C99-mEGFP in GPMVs prepared in the continuous presence of the triple inhibitor cocktail. P_ordered_ values were calculated from 3 independent experiments. The mean ± SD is reported on the graph for v vesicles.

γ-secretase activity as well as γ-secretase inhibitors are known to modulate cholesterol and lipid metabolism as well as membrane phase behavior (47,72–74). We therefore considered the possibility that the inhibitor treatments could themselves be impacting the phase preference of C99 by changing the properties of the membrane itself. To test this possibility, we asked whether other plasma membrane proteins with known phase preferences are correctly localized in ordered and disordered domains of cells where γ-secretase activity is inhibited or in cells subjected to treatment with the triple inhibitor cocktail. For these experiments, we utilized YFP-GL-GPI, a GPI-anchored protein known to localize to the ordered phase of GPMVs (49) and transferrin receptor-EGFP (TfR-EGFP), a marker of non-raft domains (75). The results of these studies showed neither DAPT nor the triple inhibitors substantially affected the *P*_*ordered*_ of YFP-GL-GPI or TfR-EGFP (**Figure 4**). Thus, even in the presence of the inhibitors, the preference of proteins and lipids for ordered and disordered domains was preserved, at least for the representative examples studied here. This implies the localization of C99-mEGFP to disordered domains is not induced by the inhibitor treatment, per se, but instead reflects its intrinsic membrane phase preference.

**Figure 4.**
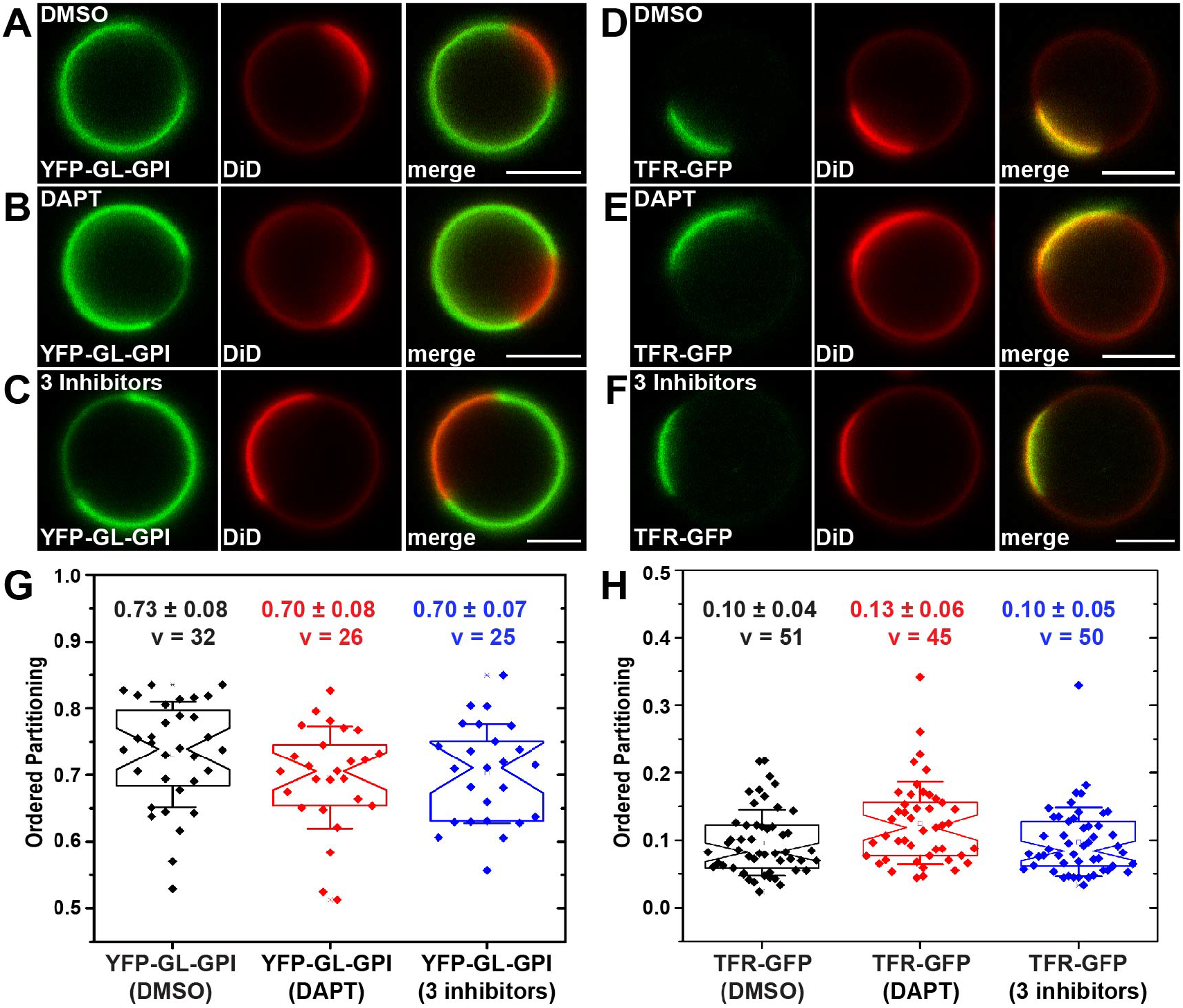
The phase preferences of the raft protein YFP-GL-GPI and non-raft protein TfR-GFP are preserved in the presence of a mixture of α-secretase, γ-secretase, and caspase inhibitors. **(A-F)** Representative images of individual GPMVs isolated from HeLa cells expressing YFP-GL-GPI (A-C) or TfR-GFP (D-F). GPMVs were prepared in the presence of either DMSO (A, D), DAPT (B, E), or a mixture of α-secretase, γ-secretase and caspase inhibitors (C, F) as described in the Materials and Methods and imaged at room temperature. Fluorescence of YFP-GL-GPI and TfR-GFP are shown in green and DiD is shown in red. Scale bars, 5 μm. **(G-H)** Ordered domain partitioning of YFP-GL-GPI (G) and TfR-GFP obtained in the presence of DMSO, DAPT, or a triple inhibitor cocktail containing α-secretase, γ-secretase, and caspase inhibitors. Ordered domain partitioning was quantified from 2-3 independent experiments with >10 GPMVs collected per replicate. Each data point represents a P_ordered_ value measured from a single GPMV. The mean ± SD for v vesicles is reported on the graph.

### Disruption of the cholesterol binding site, disruption of the dimerization interface of C99, or introduction of familial Alzheimer’s disease mutations in no case changes the preference of C99 for disordered domains

We next examined potential mechanisms that regulate the association of C99 with disordered domains. Both APP and C99 have been reported to form homodimers (76–80). Purified C99 has also been shown to interact with cholesterol (45), and the cholesterol binding site overlaps with a GXXXG motif that has an established role in dimer formation (80). In addition, multiple disease-associated mutations of full-length APP have been identified, many of which reside within these regions of the C99 protein (81–85).

To explore whether cholesterol binding or dimer formation are responsible for C99-mEGFP disordered localization in phase-separated GPMVs, we generated mutations in two sites reported to be required for cholesterol binding, I703A and E693A (45,86). APP/C99-G704 has been also been proposed to be essential for dimer formation as well as cholesterol binding (45,80). We thus also generated a G704L mutant which should disrupt both dimer formation and cholesterol binding (80). Both the cholesterol binding and dimerization mutants of C99 maintained the same disordered-phase preference as wild type C99-mEGFP (**Figure 5A-D, F**).

**Figure 5.**
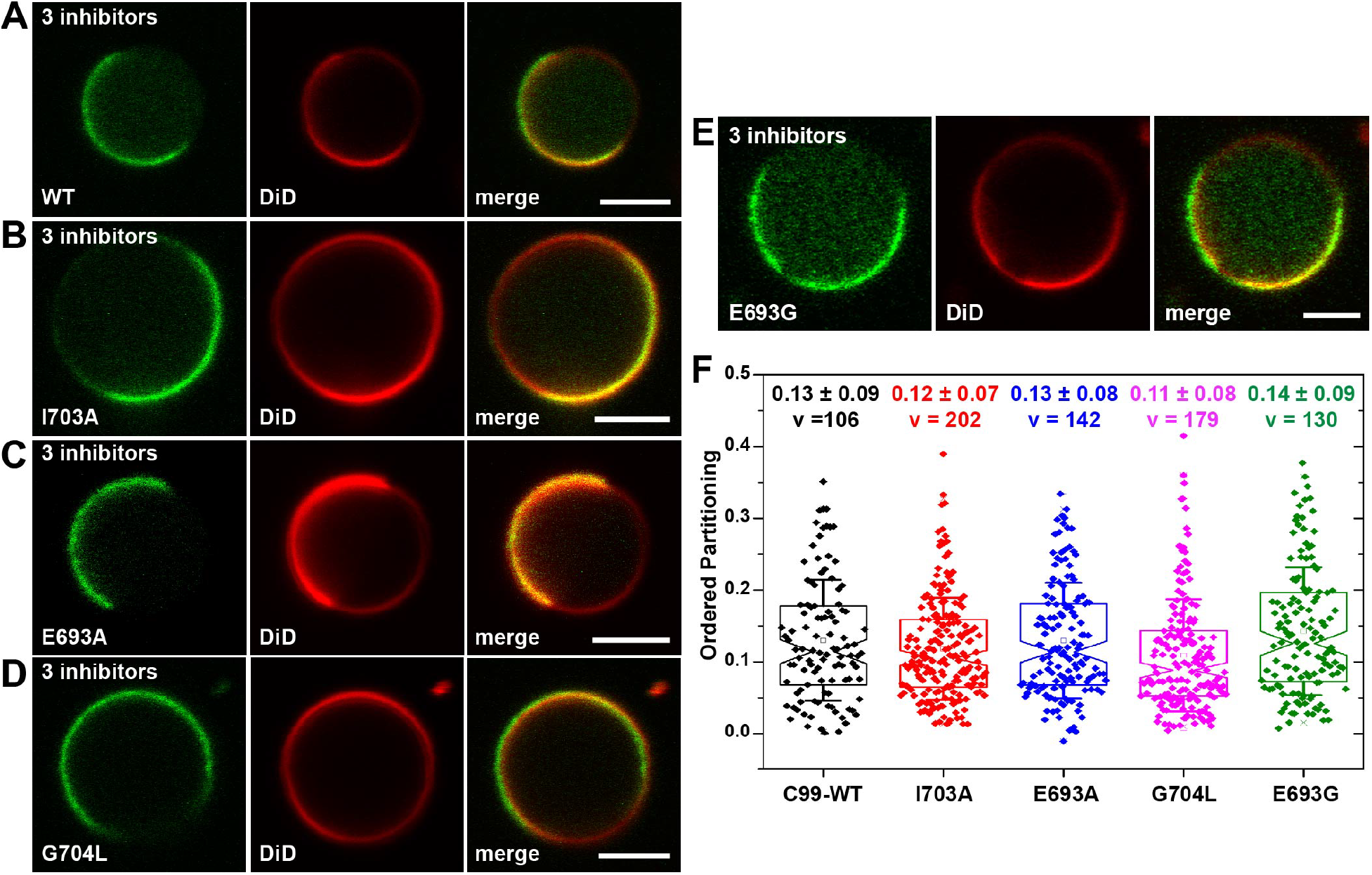
Localization of C99 in disordered domains occurs independently of its cholesterol binding activity, homodimerization, or the Alzheimer disease-associated Arctic mutation. **(A-E)** Representative examples of GPMVs derived from HeLa cells expressing either (A) WT C99-mEGFP or the following C99-mEGFP mutants: (B) I703A, a mutation reported to decrease cholesterol binding; (C) E693A, another mutation that disrupts cholesterol binding; (D) G704L, a mutation that disrupts cholesterol binding and the GXXXG dimerization motif; and (E) the familial AD Arctic E693G mutation. In A-E, GFP fluorescence is shown in green and DiD fluorescence is shown in red. Scale bars, 5 μm. **(F)** Quantification of ordered domain partitioning for WT C99-mEGFP and its mutants. P_ordered_ values were quantified as described in the Materials and Methods section and in Supporting Figure S1. For each construct, the values given in panel E represent the mean ± SD for v vesicles. The data in the graph for WT C99-mEGFP are reproduced from Figure 3. Each experiment was repeated at least three times.

We also investigated the effects of a familial Alzheimer’s disease mutation on the phase preference of C99. The site E693 is subject to multiple familial APP mutations (81–85). We chose to examine the well-studied E693G mutant (also known as Arctic mutation) (81). In GPMVs, C99-E693G-mEGFP also localized strongly to disordered domains (**Figure 5E, F).** Overall, these data show that the localization of C99-mEGFP to disordered domains occurs independently of cholesterol binding, dimer formation, or the Arctic mutation.

### C99-mEGFP localizes to disordered domains in neuronal membranes

Up to this point, all of our experiments were performed in GPMVs derived from HeLa cells. To test whether C99-mEGFP shows a similar phase preference in a more physiologically relevant cell model and lipid environment, we examined it in SH-SY5Y cells, a human neuroblastoma cell line that can be differentiated to form neurons by addition of 10μM retinoic acid (RA) (87,88). As for the case of HeLa cells, the SH-SY5Y cells were transfected with C99-mEGFP, then treated with triple inhibitor cocktail in order to prevent cleavage of C99 by γ-secretase, α-secretase, and caspases prior to and during the formation of GPMVs.

Since to our knowledge there is no prior literature that addresses whether SH-SY5Y cells form GPMVs, we first confirmed that GPMVs could indeed be generated from both undifferentiated and differentiated SH-SY5Y cells (**Figure 6A**). At room temperature, the GPMV membranes appeared to populate a uniform single phase, but phase separation was evident upon lowering the temperature into the 2°-14°C range. As shown in **Figure 6B**, in GPMVs isolated from RA-differentiated neurons, C99-mEGFP shows a variable level of EGFP on the membrane as well as luminal GFP, similar to its behavior in HeLa cells (**Figure 3**). The transfection efficiency for C99-mEGFP in both SH-SY5Y cells and neurons was low. Nonetheless, we could identify enough phase-separated GPMVs to quantify the phase preference of C99-mEGFP (**Figure 6B-D**). We obtained *P*_*ordered*_ of 0.10 ± 0.08 (v = 27) for GPMVs from undifferentiated SH-SY5Y cells and 0.14 ± 0.09 (v = 16) from neurons (Figure 6D). These values are similar to those measured in GPMVs from HeLa cells (**Figure 3C**). Thus, even in SH-SY5Y cells and differentiated neurons, a more physiologically relevant environment for C99, C99-mEGFP predominantly partitions within non-raft regions of the membrane.

**Figure 6.**
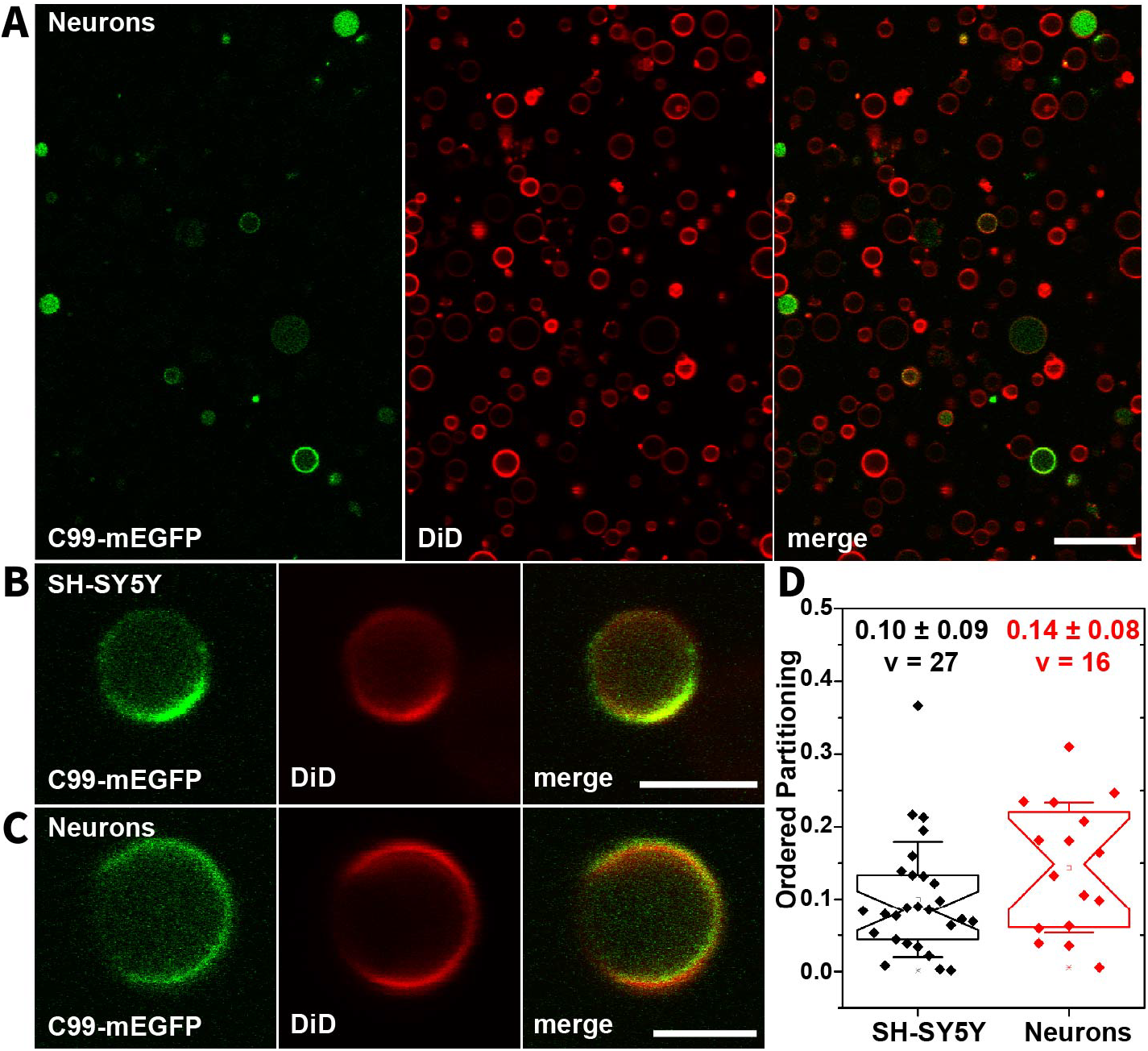
C99-mEGFP localizes in the disordered phase of GPMVs derived from SH-SY5Y cells and SH-SY5Y-derived neurons. (**A**) Representative panorama image of GPMVs generated from neuronal cells expressing C99-mEGFP. Scale bar, 20 μm. (**B, C**) C99-mEGFP shows a strict preference for the disordered phase in GPMVs generated from undifferentiated SH-SY5Y cells (B) or SH-SY5Y-derived neurons generated by retinoic acid differentiation (C) in the continuous presence of a mixture of α-secretase, γ-secretase, and caspase inhibitors. Scale bar, 5 μm. In A-C, GFP fluorescence is shown in green and DiD fluorescence is red. (**D**) Ordered domain partitioning of C99-mEGFP in GPMVs derived from undifferentiated SH-SY5Y and SH-SY5Y cells differentiated to form neurons. Images were collected at temperatures between 1-20°C for SH-SY5Y cells and 2-14°C from neurons derived from SH-SY5Y cells to increase the percentage of phase-separated vesicles available for analysis. The numbers given in panel D represent the mean ± SD for v vesicles. Each experiment was repeated three times.

## Discussion

The association of APP and its proteolytic fragments with ordered raft domains has been an area of intense research (13–18,89,90). Previous studies relying on biochemical isolation of DRMs or detergent insoluble membranes have detected the association of C-terminal fragments of APP with rafts to varying degrees (28,33). However, these approaches are known to have multiple pitfalls (19,41,42,91,92), resulting in lack of conclusive evidence for the raft association of C99. Because methods are still not available to reliably investigate the association of proteins with rafts in living cells we here visualized and quantified the affinity of C99 for raft versus non-raft domains in cell-derived GPMVs (51,52,54,62). While GPMVs do not fully recapitulate all of the features of living cell membranes, they have proven to be a valuable tool to investigate mechanisms that control the affinity of proteins for raft versus non-raft domains, and to our knowledge represent the best currently available model to address this question (49,60,61,93–96).

The major conclusion of our study is that C99-mEGFP is 85-90% excluded from raft domains in GPMVs derived from HeLa cells, SH-SY5Y cells, and neurons derived from SH-SY5Y cells. Indeed, we had difficulty even detecting the 10-15% fraction of C99 present in the ordered or raft phase of GPMVs in our studies. Since C83-mEGFP appears to be the predominant APP-derived species in cells treated with DAPT alone, our results suggest the same is true for C83-EGFP. These results are in agreement with our previous observation that C99 partitions almost exclusively to non-raft Ld domains in a more simplified synthetic lipid GUV model (46). Our results are at variance, however, from a recent report that an engineered γ-secretase substrate termed SB4 shows some affinity for ordered domains in supported lipid bilayers, as reported by atomic force microscopy (47). SB4 consists of the 76 C-terminal residues of APP and is thus shorter than C99, and also contains both an N-terminal AviTag and a C-terminal FLAG tag. Why SB4 appears to behave differently than C99 is not yet clear, but could reflect both different experimental conditions (direct insertion into supported model membranes versus biosynthetic incorporation of the protein into biological membranes in the present work) plus the fact that SB4 is essentially a truncated form of C83 and C99. Further investigation will be required to resolve this question.

In recent years, a number of mechanisms have been identified that help regulate the association of proteins with rafts. One of the most widely recognized is palmitoylation, also known as S-acylation (50,97). Several protein components of the amyloidogenic pathway are palmitoylated, including APP itself (33,34,98). Full length mature APP is palmitoylated within its N-terminal ectodomain on Cys186 and Cys187 (99). However, these APP palmitoylation sites are somewhat unusual in that they are localized to the extracellular (luminal) domain, rather than in the cytoplasmic domain of the protein where most palmitoylation sites of transmembrane proteins are found (99). Palmitoylation has been suggested to regulate APP association with DRMs and processing in rafts, as well as its dimerization (99,100). Importantly however, the C99 domain of APP lacks any cysteines and can therefore cannot undergo palmitoylation, ruling out the possibility that this form of posttranslational modification determines whether C99 itself associates with rafts.

The length and surface area of the transmembrane domain (TMD) of single-pass transmembrane proteins have also been identified as factors that help control their affinity for raft domains (50). In particular, proteins with longer TMD that also have smaller surface areas tend to exhibit higher raft affinities (50). Based simply on the sequence of its TMD, APP and C99 would be predicted to be more likely to partition within disordered regions of the membrane than in ordered raft domains (50). This is in agreement with what we observed in the experiments of this paper as well as in our previous GUV study (46). We note also a previous paper that reported simulations of a TMD-containing C99 fragment in phase-separated membranes in which the conclusion was made that C99 preferentially partitions at the interface between the Ld and Lo phases, but does not enter the Lo phase (101). This also was not observed in the our present GPMV or previous GUV studies.

Several previous experimental observations may offer clues as to why C99 shows an overwhelming preference for the disordered phase in phase-separated GUVs and cell-derived GPMVs. First, both experimental and computational studies have shown that the TMD of C99 is flexible (45,102–107). At the same time, C99 has been shown to be adaptable to changes in bilayer thickness, wherein the C-terminus of its TMD has a fixed transmembrane-end site (defined by three consecutive lysine residues) but the N-terminus of the TMD helix seems to be energetically tolerant of being either membrane-buried (in thicker bilayers) or water exposed (in thinner bilayers) (108). These observations together suggest that the TMD of C99 is both flexible and adaptable, leading it to prefer the disordered phase for entropic reasons. We also recently completed a study of the structure of C99 when it was “forced” into the raft-like environment provided by sphingomyelin and cholesterol-rich bicelles (109). In that study, C99 was seen to populate significant populations of monomer, homodimer, and homotrimer, an observation consistent with the notion that C99 is structurally and energetically frustrated in raft-like bicelles resulting a structural heterogeneity (109).

C99 also contains another structural feature that at first glance might be expected to endow the protein with a significant affinity for cholesterol-rich raft domains: a cholesterol binding site (18,45,86,110). The functional consequences of cholesterol binding by C99 are incompletely understood, but recent work suggests C99 regulates cholesterol trafficking and cellular lipid metabolism (74,111,112). Key residues in C99 involved in cholesterol binding lie in the N-terminal helix, N-terminal loop, and extracellular end of the TMD (45). While many proteins are now known to bind to cholesterol (113,114), to our knowledge the raft affinity of only a few examples of cholesterol-binding proteins have been systematically investigated. One of the best characterized examples is Perfringolysin O, a cholesterol-dependent cytolysin that exhibits intermediate affinity for rafts in its membrane inserted form (115). The multipass transmembrane protein peripheral myelin protein 22, which also contains putative cholesterol binding sites, shows a strong preference for partitioning in ordered domains in GPMVs (61). Its preference for residing within rafts, however, is retained even when both cholesterol binding sites are mutated (61). In the current study, we found that either in the presence or absence of an intact cholesterol binding site, C99 is primarily excluded from rafts. This in turn suggests binding of cholesterol per se is not sufficient to sorting of transmembrane proteins into ordered domains. We speculate that cholesterol binding could potentially even serve as a mechanism to sort single pass transmembrane proteins into disordered/non-raft domains, if it induces structural conformations that shorten the length or increase the surface area of the transmembrane region. It should also be recognized that the disordered phase domains in ordered/disordered phase-separated vesicles are cholesterol-rich—typically with cholesterol concentrations that approach that present in the ordered phase (116), such that membrane proteins with cholesterol binding sites do not need to partition into the ordered phase in order to satisfy their cholesterol binding potential.

C99 is also known to undergo homodimerization (78,80,102,117–123), a process that been reported to regulate the partitioning of some other proteins into or out of rafts (124–126). Homodimerization of C99 has clear biological consequences: for example, it decreases γ-secretase cleavage (127) and competes with cholesterol binding (80,101). We therefore asked whether the association of C99 with non-raft domains is regulated by homodimerization. We found that disruption of GFP-induced dimerization by introduction of the A206K mutation into EGFP, or mutation of a key dimer-promoting residue G704 in the TMD of C99 had no effect on the phase preference of the protein. We thus conclude that the localization of C99 to non-raft domains occurs independently of its dimerization state. Preliminary evidence also suggests the disordered preference of C99 was preserved when the C-terminal GFP tag was removed and the protein was instead labeled with an HA tag on its N-terminus (data not shown^1^), further confirming that the mEGFP moiety did not affect the localization.

Interactions of C99 with other proteins may also be important to enhance its affinity for rafts. For example, the intracellular domain of flotillin-1, itself thought to be a raft-associated protein by virtue of its association with DRMs, is thought to increase the raft association of APP (25). Notably however, flotillin has been observed to favor the disordered phase in GPMVs, a behavior that may reflect the need for cytoskeletal attachment or membrane asymmetry to help maintain its association with ordered domains in GPMVs (49). It is also important to note that, by definition, only the plasma membrane-associated pool of C99-EGFP is incorporated into GPMVs. Thus, we cannot rule out the possibility that, in intracellular membrane compartments such as endosomes or the Golgi complex, whose lipid composition differs from that of the plasma membrane, C99 may behave differently.

Finally, although the primary focus of our study was to investigate the affinity of C99-EGFP for ordered domains, we also were able to follow its proteolytic processing by directly visualizing whether GFP fluorescence was present in the lumen of the GPMVs, localized to the GPMV membrane, or both in the presence or absence of inhibitors. Consistent with the idea that C99 has a short half-life and is rapidly turned over (57,128–130), only luminal GFP fluorescence was observed in the absence of any inhibitors in our experiments. In contrast, the addition of the γ-secretase inhibitor DAPT led to the accumulation of membrane-associated GFP signal in GPMVs. This membrane-associated form of the protein was most likely C83 rather than C99, as reported by western blotting of whole cell lysates. Moreover, even in the presence of DAPT, some GFP fluorescence could be observed in the lumen of the GPMVs, leading us to probe for additional proteolytic enzymes in cell lysates. The results of these experiments suggest that C99-mEGFP is processed by both α-secretase and caspases in the presence of γ-secretase inhibitors, confirming previous reports (57,67,68,71,128,131–134). The luminal GFP signal we observed in our experiments thus likely corresponds to a mixture of AICD-mEGFP, C31-mEGFP, and GFP itself. Some GFP fluorescence could still be observed in the lumen of GPMVs even in the presence of the triple inhibitor cocktail. This indicates that additional proteolytic processing of the protein evidently occurs, although we did not pursue the identity(ies) of the putative additional proteases. In light of recent reports that GPMVs contain pores that are permeable to some hydrophilic solutes (135), further studies will be required to determine whether these soluble cleavage products are quantitatively retained in the GPMV lumen. It is also currently unclear if proteolytic processing of C99-mEGFP only occurs in unperturbed cells or continues as GPMVs are being prepared and imaged. Nevertheless, our current results raise the possibility that GPMVs may be a useful model to study processing of plasma membrane proteins that undergo proteolytic cleavage to liberate soluble cytoplasmic fragments.

In summary, our results suggest a model in which C99 localizes primarily to non-raft domains. This in turn implies either that cleavage of the protein by γ-secretase occurs predominantly in disordered regions of the membrane, or that only a minority fraction of C99 is cleaved in raft domains. Our results also cannot rule out that in living cells, C99 is preferentially localized to ordered domains via protein-protein interactions that are not maintained under conditions of GPMV preps. In future studies, it will be critical to determine whether γ-secretase does indeed associate with ordered raft domains as previously suggested from biochemical studies or instead also is operative in disordered membrane domains. This is especially important given growing evidence that γ-secretase exists in complexes with other secretases (64,136,137). Answering these questions will provide important steps forward toward fully understanding how specialized membrane nanodomains contribute to the development of Alzheimer’s disease.

### Experimental Procedures

#### Materials

The γ-secretase inhibitor DAPT was obtained from Selleckchem (cat# S2215). The α-secretase (ADAM10) specific inhibitor GI254023X was from Aobiou (cat# AOB3611). The cell permeable pan-caspase inhibitor z-VAD was purchased from UBPBio (cat# F7110). HEPES (cat# H3375) and calcium chloride (cat# C1016) were purchased from Sigma. Sodium chloride (cat# BP358-212) and paraformaldehyde (Cat#O4042-500) were procured from Fisher chemicals. EM grade paraformaldehyde solution (Cat# 15714) was from Electron Microscopy Sciences, PA. Dithiothreitol was obtained from RPI (cat# 11000-1.0). Mounting media Prolong (R) Gold antifade with DAPI Molecular probes (Cat# 8961S) was procured from Cell Signaling Technologies. Prolong Gold antifade reagent without DAPI (cat# P36930) was purchased from Invitrogen. DiD was purchased from Thermo Fisher Scientific (cat# D307).

#### Plasmids

An overview of the plasmids used in this study is provided in Table 1. The plasmid pEGFP-N1-C99-GFP, referred here as C99-EGFP, was a gift from Dr. Paola Pizzo (57). Plasmid C99-mEGFP was derived from C99-EGFP. It carries the signal peptide of the human APP gene (MLPGLALLLLAAWTARA) followed directly by the β-secretase cleavage product of the full-length human amyloid precursor protein (residues 672−770, C99). A single G-codon spacer connects to the 2^nd^ amino acid codon (Val) of EGFP. The N terminal EGFP initial Met and Kozak sequence were deleted to minimize alternative ribosomal translational initiation (138,139). To eliminate EGFP-directed dimerization, EGFP was made monomeric by substituting GFP A206K (66,140). The plasmid and its protein product are referred to as C99-mEGFP. Using C99-mEGFP as a backbone, four mutants were constructed using a Q5 site directed mutagenesis kit (New England Biolabs), corresponding to APP mutations E693A, E693G, I703A and G704L. YFP-GL-GPI was as described previously (141,142). TfR-EGFP was a gift from Dr. Ilya Levental (75).

**Table 1.** Constructs used in this study.

| **Construct** | **Identity** | **Source** |
| --- | --- | --- |
| C99-EGFP | C99 | Gift from Dr. Paola Pizzo (57) |
| C99-mEGFP | C99, full APP signal sequence included, Kozak and 1<sup>st</sup> Methionine sequence before GFP deleted, A206K mutation introduced into EGFP | This study |
| C99-E693A | Cholesterol binding mutant | This study |
| C99-I703A | Cholesterol binding mutant | This study |
| C99-G704L | Cholesterol binding and dimerization mutant | This study |
| C99-E693G | Familial Alzheimer's disease mutation (Arctic) | This study |
| YFP-GL-GPI | Model GPI-anchored protein; raft marker | (141,142). |
| TfR-GFP | Transferrin receptor; non-raft marker | Gift from Dr. Ilya Levental (75). |

#### Cell culture

All cells were maintained in a tissue culture incubator with humidified air supplemented with 5% CO2 at 37°C. HeLa (ATCC cat# CCL-2) and SHSY5Y (CRL2266 at passage 27) cells were obtained from ATCC. HeLa cells were maintained in DMEM (Gibco cat# 11885-084) supplemented with 10% fetal bovine serum (FBS, Gibco 26140-079), 1% penicillin/streptomycin (P/S), and 1% L-glutamine. SH-SY5Y cells were maintained in 1:1 DMEM/F12 with 15 mM HEPES and Glutamine (Corning cat# 10-092-CV) supplemented with 10% FBS and 1% P/S. Before passaging SH-SY5Y cells, the spent media was collected and centrifuged at 150xg for 5 minutes to collect the floating fraction (87). Passage of SH-SY5Y cells was not allowed to exceed 15 from the original P27 from ATCC.

SH-SY5Y-derived neurons were obtained from cells grown in plates pre-coated with 0.1 mg/mL Poly-D-Lysine (Gibco A3890401) to which 50,000 cells/cm^2^ were seeded and grown for 2 days to ~ 75-80% confluency. Differentiation was initiated by exchanging the SH-SY5Y media with Neuronal differentiation media (87,88). Differentiating /ed cells were only observed prior to changing to fresh media and were protected from light during handling (143). 80% of the differentiation media was changed every 2 days. The differentiation media consisted of Neurobasal medium with B27 supplement (Gibco cat# 17504-044) and 1% Glutamax (Gibco cat# 35050-061) from which an aliquot is pre-warmed and freshly supplemented each time to a final concentration of 10 μM all *trans* retinoic acid (ATRA) (MilliporeSigma, cat# PHR1187). ATRA aliquots were previously prepared in DMSO to a concentration of 3 mg/mL (10 mM) in an anaerobic chamber, aliquoted, protected from light and stored at −80°C.

#### GPMV preparation

GPMVs were prepared based on an established protocol (52) with the following modifications. Briefly, ~ 2.5-3×10^5^ HeLa cells were seeded in a 100 mm plate. 15-24 h later when the cells had reached ~ 30-35% confluency they were transfected with 3μg DNA/100 mm plate using a 3:1 ratio Fugene:DNA with C99-mEGFP plasmids in DMEM media without antibiotics or serum (base media). After 4-6 h, fresh complete media was added. 24 h after transfection, the complete media was exchanged for fresh media supplemented with either 20 μM DAPT alone, a combination of 20 μM DAPT, 20 μM GI254023X and 200 μM z-VAD, or other inhibitor combinations as indicated in the text and figure captions.

After an additional 24 h, when the cells had reached 60-70% confluency, they were rinsed twice with 7 mL C99-GPMV buffer supplemented with inhibitors (50 mM HEPES, 150 mM NaCl, 2 mM CaCl_2_, pH 7.4, with 20 μM DAPT, 20 μM GI254023X, 100 μM zVAD, with 25 mM PFA and 2 mM DTT). Cells were then incubated in 5 mL of the same buffer at 37°C with gentle shaking at 70 rpm for 1.5 h. Parallel to GPMV formation, four glass coverslips were surface treated as follows: first rinsed twice in 100% ethanol, rinsed 2-3 times in milliQ water and then incubated in 0.1% BSA dissolved in base GPMV buffer and filtered (0.2 μm). Just before use the coverslips were rinsed once in milliQ water and left to air dry.

After 1.5 h, the 5 mL of C99-GPMV containing buffer was collected, transferred to a 15 mL tube, and supplemented with 20 μM DAPT, 20 μM GI254023X, and 1.5 μg/mL DID (Ld phase marker (53,65), diluted from a 1 mg/mL stock in EtOH). The tube was gently mixed by inversion 2-3 times and decanted for 1 h at RT, after which a 260 μL aliquot of GPMVs was collected near the bottom of the tube to avoid cell debris. The aliquot was added to a sandwich made with two coverslips (Marienfeld GmbH 22 x 22mm, 1.5H, cat# 107052) pre-coated with 0.1% BSA and a 1 mm silicon spacer (EMS, cat# 70336-10), and allowed to settle for 40-60 min on a cooling stage directly on the microscope prior to imaging. Inhibitors for caspases, γ- and α-secretase were present during the entire ~36 hours including acquisition of images. The final DMSO concentration was held below 0.5%.

The same procedure was used for GPMVs generated from SH-SY5Y or neurons, with the following modifications: for SH-SY5Y cells, 4×10^6^ cells/100mm plate were seeded and transfections were performed using Lipofectamine 3000 using a 3:2 ratio of Lipofectamine 3000:DNA. For neurons, transfection was performed on the 9^th^ day with differentiation media using a 1:2 ratio of DNA:Neuromag reagent (Ozbiosciences, cat# NM50500). A total of 2μg DNA/ 60mm plates was added directly in the differentiation media, and fresh media was exchanged in after 4 h.

For control experiments, HeLa cells were transiently transfected with YFP-GL-GPI or TfR-GFP. 24 h post transfection cells were treated with either DMSO, DAPT (20 μM) or cocktail of inhibitors (20μ M DAPT, 20 μM GI254023X and 200 μM z-VAD). 24 hours after addition of DMSO or inhibitor(s), GPMVs were generated in PFA-DTT containing GPMV buffer supplemented with DMSO or inhibitor(s) using a protocol similar to that described above for C99-EGFP transfected cells. A 275 μL aliquot of GPMVs was collected, placed in between 2 coverslips separated by a 1.0 mm spacer, and imaged at room temperature using an LSM 880 confocal microscope as described below.

#### GPMV imaging

GPMVs were imaged either using a Zeiss LSM 510 confocal microscope or a Zeiss LSM 880 confocal microscope. For experiments performed using the LSM 510, a Plan-Neofluar 40x/1.30 Oil DIC objective was used. The confocal pinholes were set at 150 μm aperture for both channels in all experiments. The fluorophores were excited using the 488 nm line of a 40 mW Argon laser set at no more than 5-10% power (reduced at source to 50%, for final 2.5-5%) for EGFP signal, and the 633 nm line of a HeNe laser set at no more than 2% power for DiD. Images were collected at 1-2x zoom for sample overview or 10X digital zoom for quantifying phase partition localization. The stage was cooled using a TS-4 MP from Physitemp Instruments (Clifton, NJ, USA) with a custom-fitted adapter for the LSM-510 stage. The cooling stage was aided by connecting the lines to a refrigerated pump (Neslab Instruments Inc. model RTE-111) set at 10°C. Images were collected over a nominal temperature range of 4-22°C for GPMVs from HeLa cells, 1-20°C from SH-SY5Y cells and 2-14°C from neurons derived from SH-SY5Y cells. Because an oil objective in direct contact with the coverslip was used for these experiments, these temperatures are only approximate.

Imaging of GPMVs containing YFP-GL-GPI and TfR-GFP, and data for Figure 1 for C99-EGFP and C99-mEGFP were collected on an LSM 880 confocal microscope using C-Apochromat 40x/1.2 W Korr FCS M27 objective at room temperature. The confocal pinhole was set to 150 μm aperture for all channels. Single Z plane images of individual GPMVs at 10x optical zoom were acquired using the 488 nm laser line for GFP or YFP laser lines and 633 nm line for DiD. For imaging GPMVs containing YFP-GL-GPI, laser output was set to no more than 2.0%. Laser output for GPMVs with TFR-GFP not more than 6.0% for 488 nm and for 633 nm laser output was no more than 2%. For presentation purposes, images were processed using ImageJ software.

#### Quantification of raft association

Quantification of ordered domain partitioning for DiD, YFP-GL-GPI, and TfR-GFP was carried out essentially as described previously (60,61,94). In brief, a line scan was drawn across the GPMV using Image J to determine the fluorescence intensity in both the ordered and disordered phases. A moving average of 5 pixels was used to smooth the data. The ordered phase partitioning fraction P_ordered_ was then calculated using the fluorescence intensity in the ordered phase (I_ordered_) and fluorescence intensity in the disordered phase (I_disordered_) as (52,53)

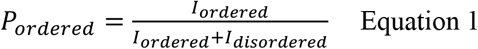

In GPMVs containing C99-EGFP and C99-mEGFP, GFP fluorescence in the ordered phase was typically extremely low, and individual GPMVs contained variable amounts of GFP fluorescence in their lumen. We therefore modified the analysis of partition coefficients as follows (**Supporting Figure S2**). Using the Plot Profile function of ImageJ, a line was drawn bisecting each GPMV at a position that sampled both the ordered and disordered domains. The line was extended into surrounding background on either side of each GPMV. Using this approach, the position of the peak fluorescence intensity was readily visualized for DiD in both the disordered phase and the ordered phase, as well as for C99-EGFP in the disordered phase. The fluorescence intensity was averaged across the three pixels at the maximum of each peak and recorded. The mean fluorescence intensity of C99-EGFP or C99-mEGFP in the ordered phase was measured at the position of the three pixels corresponding to the peak DiD fluorescence in the ordered phase. Background fluorescence was determined by averaging across 20-25 pixels at each end of the line for each channel. The ordered phase partitioning fraction coefficient was then calculated using background-subtracted values for I_ordered_ and I_disordered_ using the equation above. We also confirmed the ordered phase partitioning fraction measurements for a subset of GPMVs containing C99-mEGFP using a second method based on Azimuthal averaging as described in **Supporting Figure S5**.

#### Fluorescence microscopy of fixed cells

HeLa cells grown on glass coverslips were transfected with C99-EGFP or C99-mEGFP as described above. 24 hours post transfection, cells were treated with DMSO, DAPT (20 μM), or a triple inhibitor cocktail consisting of 20 μM DAPT (γ-secretase inhibitor), 20 μM GI254023X (α-secretase inhibitor) and 200 μM z-VAD (pan caspase inhibitor). After incubating the cells an additional 24 h with indicated inhibitors, the cells were fixed at 37° C for 15 min in 4% PFA in DPBS. The coverslips were mounted on a glass slide using Prolong Gold antifade reagent containing DAPI for nuclear staining.

Fluorescence imaging of fixed cells was performed using a Zeiss LSM 880 confocal microscope (Carl Zeiss Microscopy, Inc.; Thornwood, NY) using a Plan-Apochromat 63X/1.4 Oil DIC M20 Zeiss oil immersion objective. Fluorescence was excited using a 405 diode and the 448 nm line of an Argon/2 30mW laser and fluorescence images were collected using filter sets provided by the manufacturer. For presentation purposes, images were processed using ImageJ software.

#### Western blotting

HeLa cells were plated, transfected, and treated with inhibitors exactly as described above for GPMV preparation and were 70-80% confluent on the day of the experiment. They were placed on ice, rinsed once with 5 mL ice-cold DPBS containing 200 μM z-VAD, 20 μM DAPT, and 20 μM GI254023SX, and then lysed in 300-450 μL Lysis buffer (based on DPBS: 136.9 mM NaCl, 2.67 mM KCl, 1.47 mM KH_2_PO_4_, 8.10 mM Na_2_HPO_4_, supplemented with 50 mM Hepes pH 7.4, 1% (w/v), Nonidet P-40 (NP-40), 1% Triton X-100, 0.5% (w/v) SDS, 2 mM EDTA, and protease inhibitors 100 μL Sigma-PI8340, 100 μM DAPT, 100 μM GI254023X, 300 μM z-VAD, 200μM Pepstatin A, and just before use 2 mM PMSF). Lysis was performed on ice for 5 minutes. Next, extracts were harvested on ice using a cell scraper into 1.5 mL tubes and kept on ice, further mixed by pipetting and incubated on ice for an additional 30 min. Samples were clarified by centrifugation at 20,000 x g for 30 min at 4°C. Supernatant and aliquots for protein quantification were collected into fresh tubes kept on ice, flash frozen in liquid N_2_ and stored in −80°C. Protein concentration was determined by Bradford assay (Pierce cat# 1863028) using a linearized Bradford protocol (144). Western blots were performed using 10 μg/lane of extracts, and run on 10% Bis-Tris acrylamide precast gels (NuPAGE cat# NP0301) with MES as running buffer. For some experiments, MOPS was used as a running buffer instead of MES to increase resolution of bands between 28-38kDa, at the expense of resolving lower molecular weights.

To detect the N-terminus of C99/Aβ, blots were probed with mouse IgG mAb 82E1 (IBL cat# 10323; lot 1D-421) used at 0.5 μg/mL. The C-terminal domain of C99 was detected using a 1:1000 dilution of APP-CT20, a rabbit polyclonal raised against the last 20 amino acids of APP 751-770 (Sigma-Aldrich cat# 17610). GFP was probed using mouse mAb JL-8 (dilution 1:5000; Takara, cat# 632381) or rabbit mAb GFP (D5.1) XP (dilution 1:1000; Cell Signaling, cat# 2956).

To detect actin, blots were probed with mouse IgG1 mAb MCA-5J11 against all actin isoforms (1:1000 dilution, Encorbio, cat# MCA-5J11).

Protein signal was detected using ECL substrate (Pierce #32209) incubated at room temperature for 5 min with gentle hand mixing. Images were acquired using the Amersham Imager 600 (GE) set on chemiluminescence, with high dynamic rage and colorimetric marker options selected.

#### Statistical analysis

Measurements of the ordered domain partitioning fraction of C99-EGFP and C99-mEGFP were obtained from 2 or more independent experiments and from at least 2 independent experiments for YFP-GL-GPI and TfR-GFP. P_ordered_ values were plotted as notched boxes with error bars showing one standard deviation (SD). Each data point represents a measurement from a single GPMV. The actual numerical values are also reported as mean ± SD across all independent experiments for each protein and treatment.

## Data Availability

All data either are presented in the article, are in the supporting information, or are available from the corresponding author (Anne K. Kenworthy) upon request.

## Acknowledgements

We thank Krishnan Raghunathan and Nico Fricke for assistance at early stages of this study, the Vanderbilt Cell Imaging Shared Resource (CISR) for access to confocal microscopes and Sean Schaffer for assistance optimizing conditions to image GPMV samples.

## Author Contributions

AT and RC data curation; RC, AT, YP, and JMH, formal analysis; RC and AT investigation; AT, RC, and AKK writing-original draft; RC, AT, YP, AH, JMH, AKK, and CRS write— review and editing; AT, RC, and JMH, methodology; AKK, AT, RC, and CRS conceptualization; YP and AH resources; AKK and CRS supervision; AKK and CRS project administration.

## Funding and Additional Information

The CISR is supported by NIH grants CA68485, DK20593, DK58404, DK59637, and EY08126. This work was supported by NIH 1RF1 AG056147 to AKK and CRS. JMH was supported by NIH T32 CA00958229 and by F31 AG061984. The content is solely the responsibility of the authors and does not necessarily represent the official views of the National Institutes of Health.

## Abbreviations

Aβ: amyloid-β
APP: amyloid precursor protein
BACE1: β-site amyloid precursor protein cleaving enzyme 1
DRM: detergent resistant membrane
GPMV: giant plasma membrane vesicle
GUV: giant unilamellar vesicle
Ld: liquid disordered
Lo: liquid ordered
TfR: transferrin receptor
TMD: transmembrane domain
v: number of GPMVs measured

## Footnotes

1. A. Tiwari and R. Capone, unpublished observations

