## Supporting information for "The C99 domain of the amyloid precursor protein is a disordered membrane phase-preferring protein"

Supporting information includes:

Figure S1

Figure S2

Figure S3

Figure S4

Figure S5

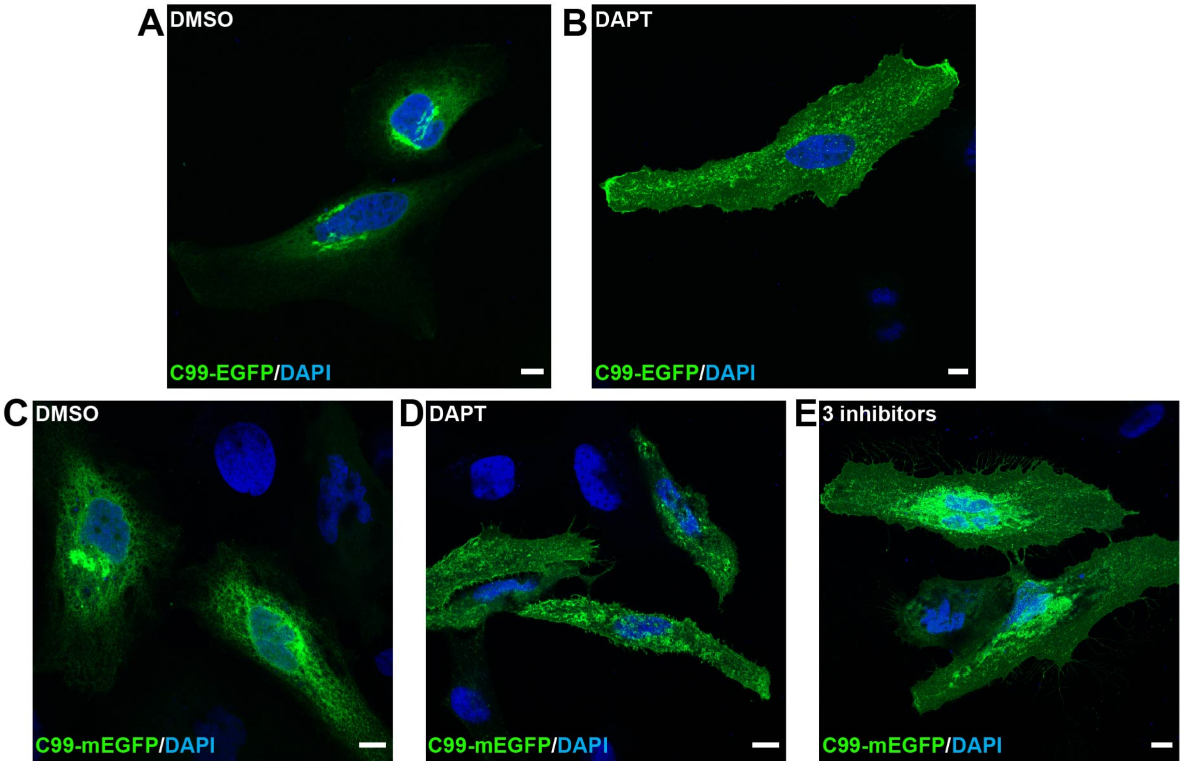


**Figure S1. C99-EGFP and C99-mEGFP localize to the plasma membrane following treatment of HeLa cells with a γ-secretase inhibitor or a mixture of inhibitors against α-secretase, γ-secretase, and pan-caspase inhibitor.** HeLa cells were seeded on glass coverslips and transiently transfected with C99-EGFP (**A, B**) or C99-mEGFP (**C, D, E**). They were then treated with DMSO (A, C), 20µM DAPT (B, D), or mixture of α-secretase inhibitor (GI254023X, 20µM), γ-secretase (DAPT, 20µM) and pan-caspase inhibitor (z-VAD 200µM) (E). 24 h after treatment, the cells were fixed, stained with DAPI, mounted and imaged by confocal microscopy. GFP fluorescence is shown in green and DAPI staining of nuclei is shown in blue. Scale bars, 10 µm.

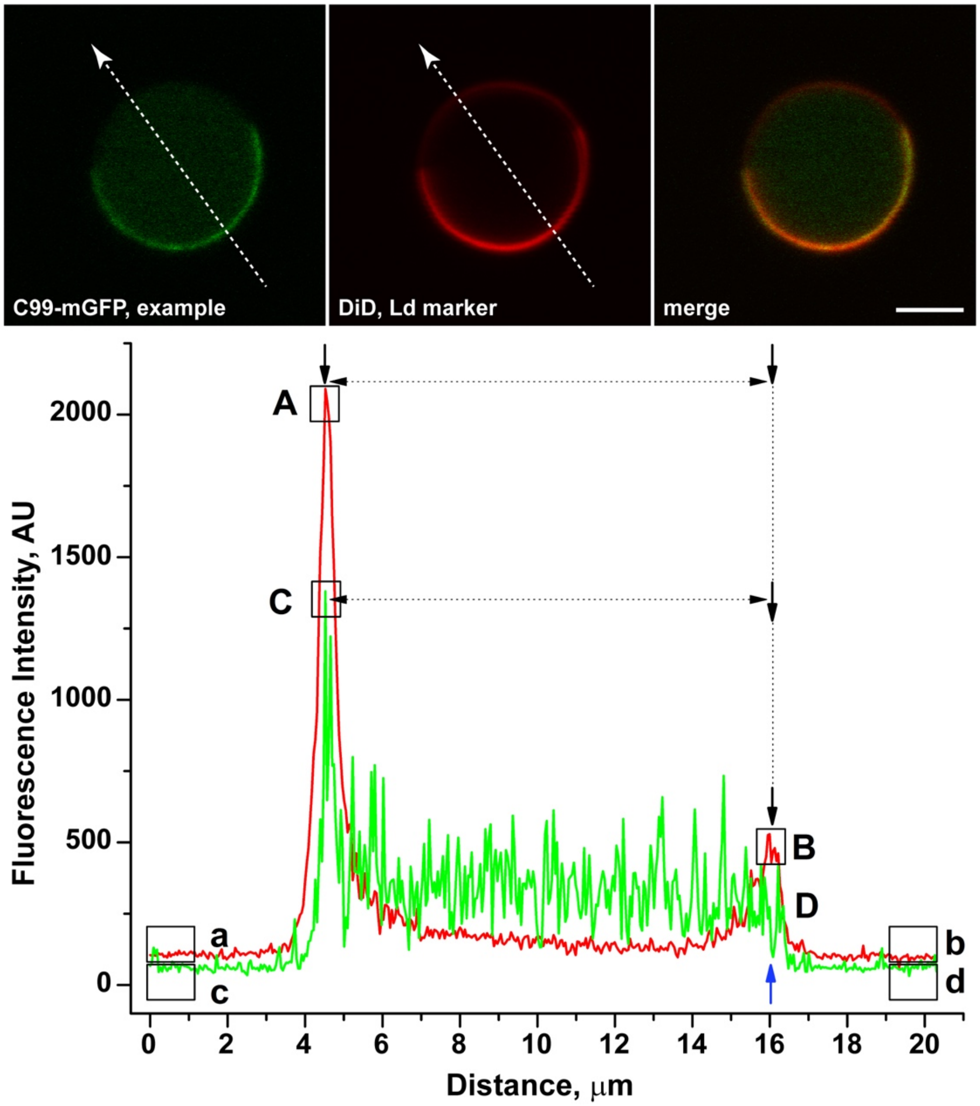


**Figure S2. Method used to analyze ordered domain partitioning for C99-EGFP and C99-mEGFP constructs in GPMVs.** The top panel shows an example of a GPMV and the positioning of the line used to analyze the fluorescence intensity. The bottom panel shows a graph of fluorescence intensity across the line in both the green channel (GFP) and red channel (DiD). Peak A marks the fluorescence intensity for DiD in the disordered phase and Peak B is the fluorescence intensity for DiD in the ordered phase. Peak C corresponds to the fluorescence intensity of C99 in the disordered phase. The expected position of peak D (blue arrow) was inferred from the position of peak B and was used as a measure of the fluorescence intensity of C99 in the ordered phase. For quantitation purposes, the fluorescence intensity was averaged across the three pixels at the maximum of each peak (boxes in figure). Background fluorescence was determined by averaging across 20-25 pixels at each end of the line for each channel (a and b for DiD, and c and d for C99-EGFP or mEGFP). Scale bar, 5 μm.

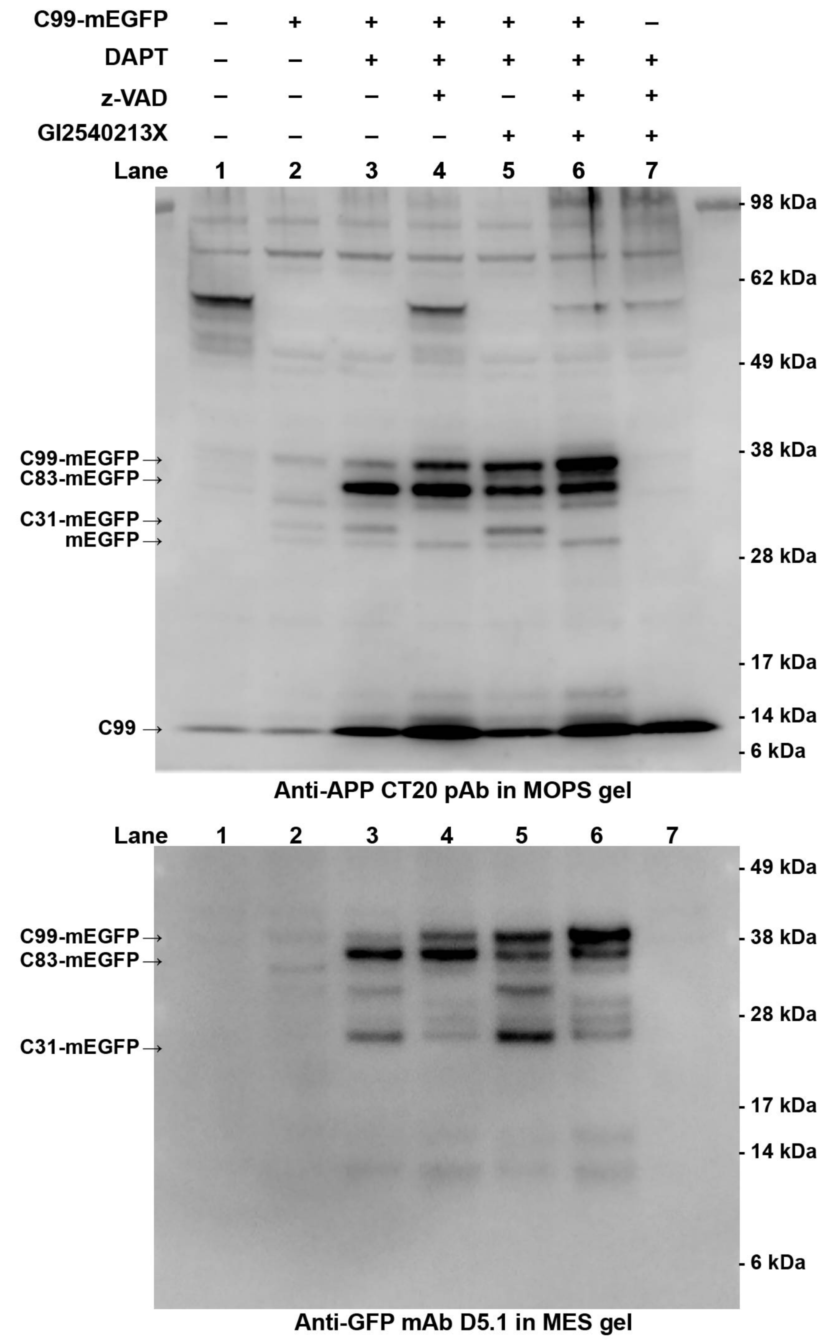


**Figure S3. Examples of Western blots probed with additional APP and GFP antibodies.** The protein extracts shown in Figure 2 were probed in the top panel with anti-APP polyclonal antibody CT20 which recognizes the last 20 amino acids of APP and therefore also C99, C83 and other C-terminal containing fragments. In the lower panel, another anti-GFP antibody was used where a smaller molecular weight fragment can be observed. Overall, a trend similar is observed to that presented in Figure 2, i.e., the levels of overexpressed C99-mEGFP are increased in the presence of the three inhibitors.

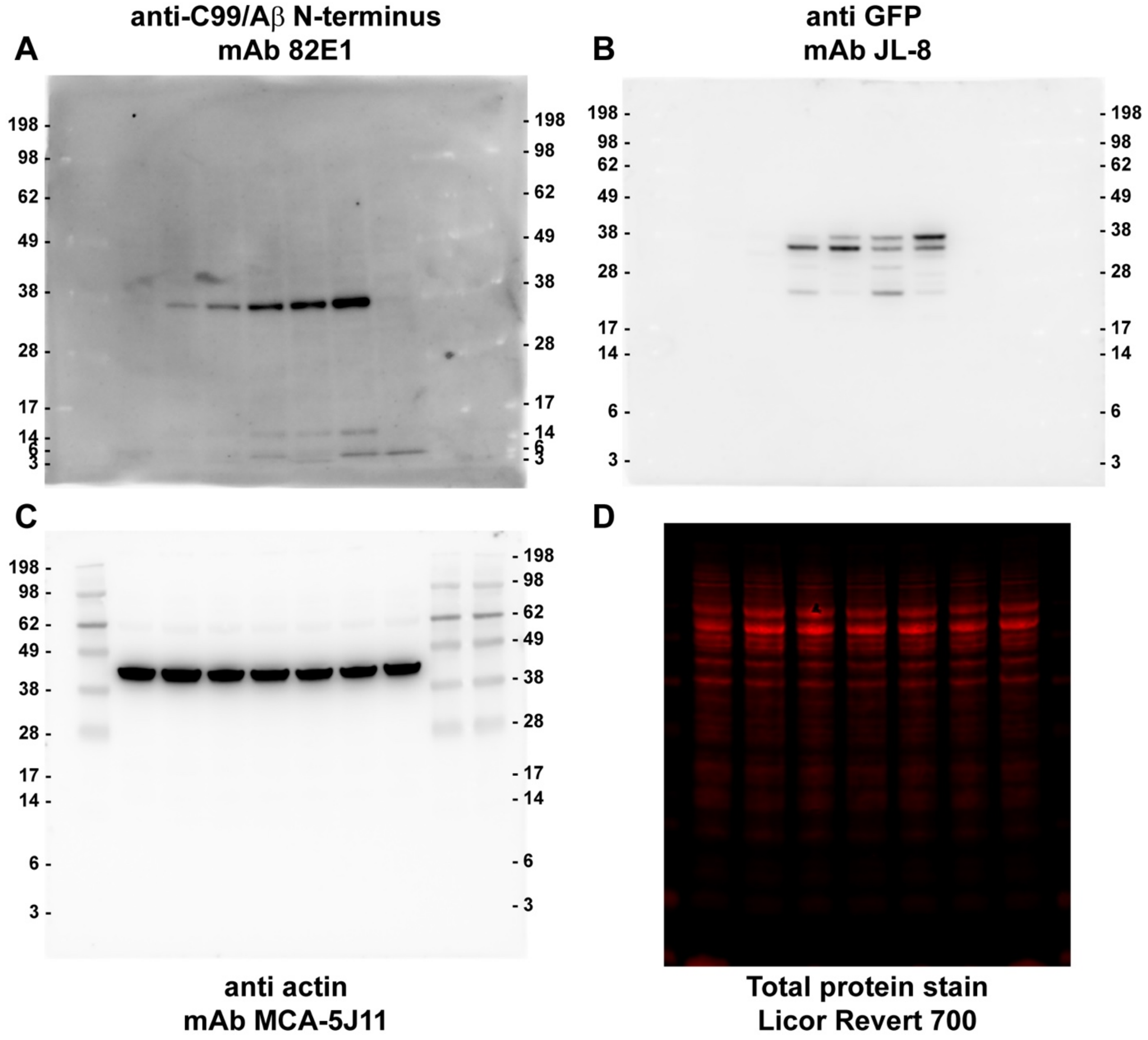


**Figure S4. Complete nitrocellulose membrane blots used to generate Figure 2.** (**A**) anti-C99 mAb 82E1; (**B**) anti-GFP mAb JL-8, and (**C**) anti-actin MCA-5J11. (**D**) An example of total protein and its transfer efficiency to nitrocellulose is shown using the Licor Revert 700 total protein staining.

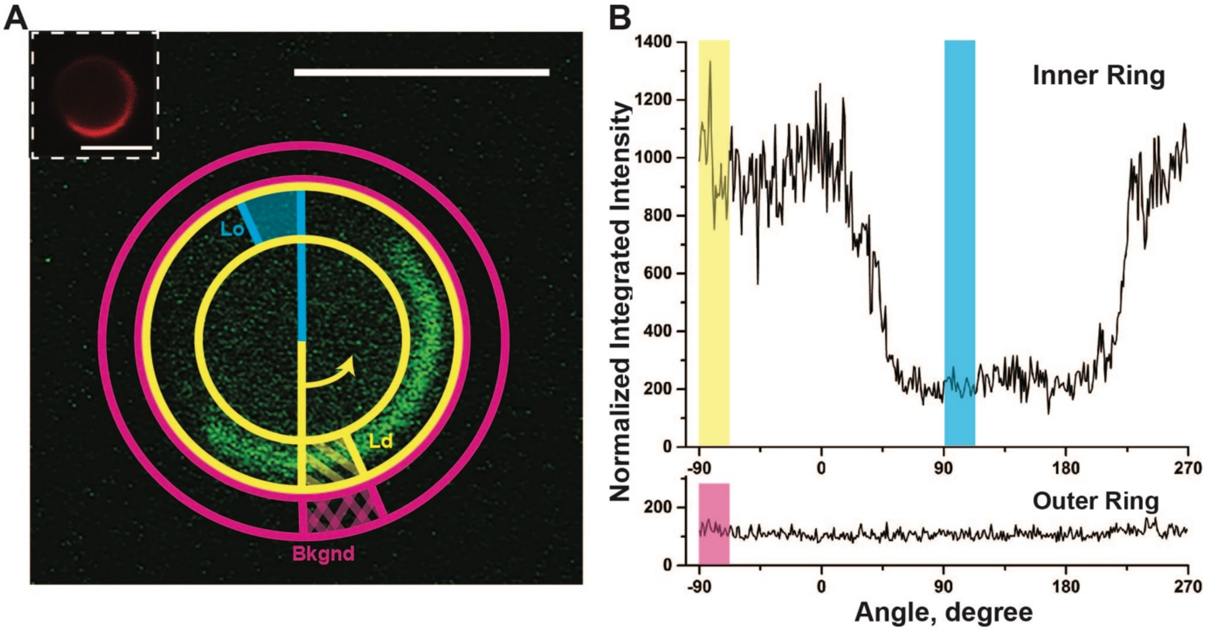


**C**

| | No. GPMV analyzed | Partition coefficient | Standard deviation |
|---|---|---|---|
| **Azimuthal Method** | 11* | 0.14 | 0.08 |
| **Line Scan Method** | 11* | 0.15 | 0.08 |
| **Line Scan Method** | 106 | 0.13 | 0.08 |

**** direct comparison of same GPMVs***

**Figure S5. Azimuthal averaging method used to validate WT C99-mEGFP ordered domain partitioning in GPMVs.** In this approach, radial integration was used to perform fluorescence signal averaging. An example of a GPMV containing C99-mEGFP in the membrane that was analyzed using this approach is shown in panel **A**. The inset shows the DiD labeling in the same GPMV. To carry out the analysis, the Azimuthal Average plugin for ImageJ by Philippe Carl (https://imagej.nih.gov/ij/plugins/azimuthal-average.html) was used to define a ring centered on the GPMV membrane (shown in yellow) or directly outside the GPMV to measure the background (shown in pink). Radial integration was performed using the plugin in a counter-clockwise direction using 360 bins to yield plots of normalized integrated intensity as a function of angle for the GPMV membrane and background (**B**). The mean intensity was measured for a small portion of the ring in the ordered domain (highlighted in blue), disordered domain (highlighted in yellow), and background (highlighted in pink). The ordered phase partitioning was then calculated as usual with Equation 1. (**C)** The mean ordered domain partitioning for C99-mEGFP obtained for a subset of GPMVs analyzed using this Azimuthal averaging method was essentially identical to that obtained using the line scan method (Supporting Figure S2). Thus, the line scan method, which was less labor intensive, was used for all analyses reported in the main body of the manuscript. Scale bars, 5 μm.
